# Multimodal Influences on Learning Walks in Desert Ants (*Cataglyphis fortis*)

**DOI:** 10.1101/2020.04.17.046839

**Authors:** Jose Adrian Vega Vermehren, Cornelia Buehlmann, Ana Sofia David Fernandes, Paul Graham

**Affiliations:** Westfaelische Hochschule, Bocholt, Germany; University of Sussex, Brighton, UK

**Keywords:** Learning walks, ant navigation, multimodal navigation, visual cues, wind

## Abstract

Ants are excellent navigators taking into account multimodal sensory information as they move through the world. To be able to accurately localise the nest at the end of a foraging journey, visual cues, wind direction and also olfactory cues need to be learnt. Learning walks are performed at the start of an ant’s foraging career or when the appearance of the nest surrounding has changed. We investigated here whether the structure of such learning walks in the desert ant *Cataglyphis fortis* takes into account wind direction in conjunction with the learning of new visual information. Ants learnt to travel back and forth between their nest and a feeder, and we then introduced a black cylinder near their nest to induce learning walks in regular foragers. By doing this across days with different prevailing wind directions, we were able to probe how ants balance the influence of different sensory modalities. We found that (i) the ants’ outwards headings are influenced by the direction of the wind with their routes deflected in such a way that they will arrive downwind of their nest when homing, (ii) a novel object along the route induces learning walks in experienced ants and (iii) the structure of learning walks is shaped by the wind direction rather than the position of the visual cue.

## Introduction

Social insect foragers are expert navigators, using a combination of innate navigational strategies and learnt information from their environment (Knaden and Graham, 2016). Early in their foraging life, ants have no information about their surroundings, and depend on idiothetic information which, via path integration (PI), is used to keep track of their approximate location relative to the nest (Mueller and Wehner, 1988). PI allows ants to explore the world safely as navigationally useful visual (Zeil, 2012; Collett et al., 2013) and olfactory (Steck, 2012; Buehlmann et al., 2015) information is learnt to increase the reliability of localising the nest or following a foraging route.

The importance of learning environmental information for a forager is demonstrated by coordinated behavioural and physiological adaptations that mark the start of an ant’s foraging career (reviewed in (Roessler, 2019)). New foragers perform a specific set of exploratory walks that allow them to systematically inspect the surroundings of their nest (reviewed in (Collett and Zeil, 2018; Zeil and Fleischmann, 2019)). During this early learning phase, ants do not search for food and only leave the nest for short periods, covering no more than a few centimetres before turning back (Wehner et al., 2004). The distance, time and path straightness then increase with each subsequent walk (Wehner et al., 2004; Fleischmann et al., 2016). Synchronised with this structured exploration, the brains of foragers show neuronal changes in the key brain areas associated with navigational systems and learning (Kuhn-Buehlmann and Wehner, 2006; Stieb et al., 2010; Stieb et al., 2012; Schmitt et al., 2016; Grob et al., 2017).

The structure of the learning walks of many species involves conspicuous turns and loops (*Cataglyphis fortis*: (Stieb et al., 2012; Fleischmann et al., 2016); *Cataglyphis bicolor:* (Wehner et al., 2004); *Ocymyrmex robustior:* (Mueller and Wehner, 2010), *Cataglyphis noda:* (Fleischmann et al., 2017), *Myrmecia croslandi:* (Jayatilaka et al., 2018); *Formica rufa:* (Nicholson et al., 1999)). Some species show fine-grained motor motifs such as voltes and pirouettes, which are small loops and nest-focussed inspections respectively, on top of the coarse structure of loops and turns (Mueller and Wehner, 2010; Fleischmann et al., 2017). Furthermore, the modulation of walking speed is also strongly correlated with the learning process of novel locations in *C. fortis* desert ants (Buehlmann et al., 2018). Thus, overall we can identify a series of changes to walking patterns which constitute an active learning process during the learning walk behaviour in a variety of ant species.

Ants do not only perform learning walks when leaving the nest for the first time but engage in learning manoeuvres under other circumstances as well. Learning walks often occur on the first journey of each day, independently of the amount of previous experience the ant has had (Graham and Collett, 2006) and also if the nest surroundings are changed significantly. For instance, if a cylinder is added near the ant’s goal location, ants engage in learning walks to update their memories of the changed environment (Nicholson et al., 1999; Mueller and Wehner, 2010).

The control of learning walks is inherently multimodal. Ants appear to use magnetic cues during early learning walks to control their orientation, with this early phase of learning walks giving an opportunity for ants to learn the configuration of celestial cues that are specific to the exact location and time of year (Fleischmann et al., 2018). However, it is less certain how learning walks are influenced by the need to learn multimodal information. Visual cues are clearly important, e.g. we see that the addition of extra visual information around the nest triggers new learning walks (Nicholson et al., 1999; Mueller and Wehner, 2010), and those species that inhabit visually rich environments invest more in the fine-control of motor motifs during learning walks (Fleischmann et al., 2017). What about the learning of information from other modalities? We know that during navigation, desert ants use information from olfaction (Buehlmann et al., 2012; Buehlmann et al., 2014, 2015), wind direction (Wehner and Duelli, 1971; Mueller and Wehner, 2007), tactile cues (Seidl and Wehner, 2006) and other sensory modalities. Wind direction is particularly interesting, as it can be used as a compass cue if it remains relatively constant over a period of time (Mueller and Wehner, 2007) and it can also be a useful carrier of olfactory information (Wolf and Wehner, 2000; Buehlmann et al., 2012; Steck, 2012; Buehlmann et al., 2014, 2015).

Here we investigate whether the structure of learning walks of *Cataglyphis fortis* desert ants takes into account wind direction in conjunction with the learning of new visual information. We trained foragers to visit a regular feeder and then introduced a black cylinder near their nest, thus learning walks were prompted in regular foragers. By doing this across days with different prevailing wind directions, we were able to probe how ants balance the influence of different modalities.

## Materials and Methods

### Ants and field site

Experiments were carried out between July and August of 2015 in the relatively featureless Tunisian salt pan near the village of Menzel Chaker (34.954897 N, 10.410396 E). Ten different nests of the desert ant species *Cataglyphis fortis* (Fig. 1A) were selected and each nest was only used once.

**Figure 1:**
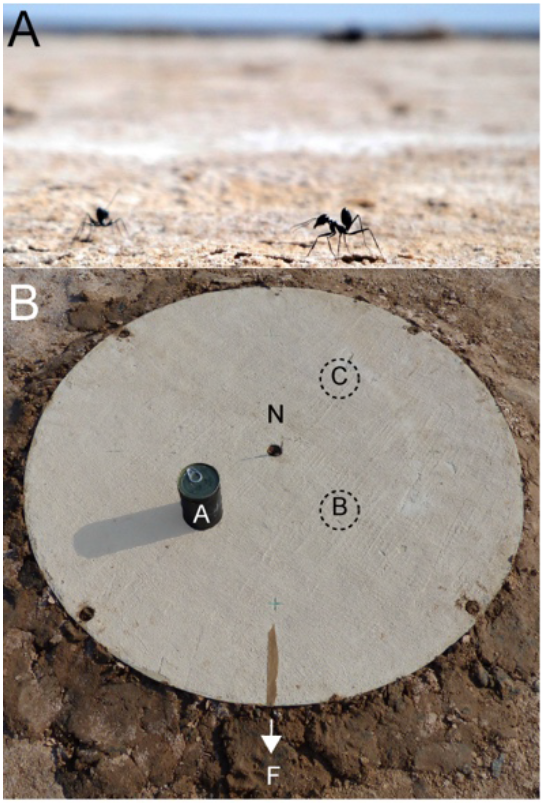
Experimental setup. **A**: Experiments were performed with the desert ant *Cataglyphis fortis* in the Tunisian salt pan. **B**: A circular arena (1 m in diameter) with a central opening (3 cm in diameter) was placed on the top of the ants’ nest (N). Ants that could leave and enter the nest only through the central hole of the arena were trained to a feeder (F) 5 m away from the nest. In tests, a black cylinder (7.5 cm in diameter, 11 cm high) was introduced 20 cm away from the nest at position A, B or C, respectively.

### Experimental procedures

A circular arena (1 m in diameter) with a central, circular opening (3 cm) was placed over the nest entrance such that outgoing ants could only leave the nest by using the central hole and crossing the arena (Fig. 1B). Ants learnt to find a feeder that was placed 5 m away from the nest providing biscuit crumbs ad libitum for at least half a day. To control the movements of ants, fluon on the walls of the feeder stopped ants climbing out, unless a wooden stick was present as a bridge, which during training, allowed ants to travel back and forth between the nest and the feeder.

The procedures for the Control recordings were to (i) remove the wooden stick at the feeder to collect a batch of foragers, (ii) put the stick back to let all ants return back to the nest, (iii) start recording departing outwards ants once most ants had entered the nest and (iv) let them walk back and forth between the nest and feeder again. Similarly, the same procedures were followed for the recordings of the learning walks that were induced by adding a novel cylinder once most ants had entered the nest (between (ii) and (iii)). Therefore, with a batch of regular foragers back in the nest, we could add the cylinder to induce learning walks on the ants’ next emergence from the nest (Cylinder condition). The black cylinder (7.5 cm in diameter, 11 cm high) was placed 20 cm away from the nest entrance in one of 3 possible locations (A, B or C) around the nest entrance (Fig. 1B). When time allowed, we collected the outwards ants in the feeder after their first exposure to the cylinder and recorded their subsequent inbound run in the presence of the novel cylinder.

Ants crossing the arena were recorded with a Panasonic DMC-F2200 high-speed camera (200 fps). A 5 cm high stick with a string attached on the upper end was placed at the centre of the arena to record the wind direction throughout the experiments. To extract wind direction, movies were viewed in VLC movie player (version 3.0.3) and the direction of the string was categorized by eye into one of 16 sectors (sector width 22.5°) every 240 frames (1.2 s). On all testing days, wind direction did not change more than 30° during an experiment. In eight out of ten days, the prevailing wind direction was 55° positive to the feeder direction (+55°). Because of the winds’ consistency, these 8 days were grouped (Jul-07 to Jul-28). Mean wind direction on Jul-06 was 31° negative (−31°) and on Aug-02 125° negative (−125°) to the feeder direction.

### Data processing and analysis

The ants’ paths were manually extracted from videos at 20 fps using Graph Click (Arizona-Software, version 3.0). Tracking started when the last shadow cast by the experimenter was out of sight. Every ant leaving the nest entrance and venturing more than 2 cm away from the nest was tracked. Ants interacting with other ants were tracked as far as possible. If no distinction between two interacting ants was possible, the files were labelled, and each ant was matched with one of the departing ants. Ants disappearing behind the cylinder (from the camera’s perspective) were followed to the edge and then matched with a departing ant (from behind the cylinder) as accurately as possible. To ensure that all inwards ants came from our feeder, we only considered homing ants that entered the arena from the feeder direction +/- 90°.

Calibration marks on the arena were used for calibration and digitised paths were smoothed with a mean filter and a window size of 3 frames. Path straightness was calculated as the quotient of the beeline, i.e. the euclidean distance between the first and last point of a path, and the actual path length (Index of straightness, IS), with paths containing gaps shifted to close gaps. For more detailed path analysis, paths were divided into chunks. The length of these chunks was set to 2 cm and the direction of each chunk and the ant’s speed in that chunk was calculated. Chunks were considered to be aligned to the nest when the angular difference to the nest was smaller than 22.5°. To find chunks with turns, we looked for chunks where the distance from the nest entrance to a chunk had reduced by more than 1 cm (half a chunk) since the previous chunk. For the analysis of walking speed, a threshold of 0.06 m/sec was set to identify path segments with low walking speed that may indicate learning (Buehlmann et al., 2018).

All data was analysed and plotted using Matlab R2017b. Statistical analysis was performed using the Statistics and Machine Learning Toolbox, as well as the Circular Statistics Toolbox (Berens, 2009). Furthermore, additional circular statistics (Batschelet, 1981) were run using Oriana 4.02 (Kovach computing services) and PAST (Hammer et al., 2001).

## Results

### Ants’ outwards headings are influenced by the direction of the wind

The Control ants’ heading direction on days with the prevailing wind (+55°; Jul-07 to Jul-28) and on days with wind 31° negative to the feeder direction (−31°) could be predicted by both the feeder and the wind direction at three different distances from the nest entrance (prevailing wind: V test, n = 216 ants, r_1_ = 10 cm, r_2_ = 25 cm, r_3_ = 45 cm, Feeder: p_1-3_ < 0.001, Mean wind: p_1-3_ < 0.001; wind −31°: V test, n = 23 ants, Feeder: p_1-3_ < 0.001, Mean wind: p_1-3_ < 0.001). The wind on Aug-02 blew in a direction almost opposite to the feeder (−125° to the feeder direction). Here, the ants’ heading direction could be predicted by the feeder but not the wind direction (V test, n = 36 ants, Feeder: p_1-3_ < 0.01, Mean wind: p_1-3_ > 0.05). Moreover, the spread in the heading directions was significantly higher than in the other two wind conditions (k test with Bonferroni correction, −30° vs. +55°, p_1-3_ > 0.05; −30° vs. −125°, p_1-3_ < 0.01; +55° vs. −125°, p_1-3_ < 0.001). Taken together, wind direction had an influence on the ants’ headings, and when in a directional conflict with the feeder direction, the ants’ directional scatter in the path heading increased significantly.

### Novel object along the route induces learning walks in experienced ants

Experienced ants navigating to the feeder were introduced to a black cylinder placed at different positions relative to the nest and learnt route respectively (Fig. 1B). To analyse the impact of the different cylinder positions on the ants’ paths but discard the influence of the wind, only experiments from days with prevailing wind were analysed here (+55°; Jul-07 to Jul-28). In general, the ants’ headings were directed (Rayleigh test, all p < 0.001) and the mean heading direction in the Control and Cylinder condition did not differ from each other (ants’ headings at r_1_ = 10 cm, r_2_ = 25 cm, r_3_ = 45 cm; Control vs Cylinder, Watson Williams tests, p < 0.05 for r_1_ in A, everything else p > 0.05; see Fig. 2). However, the spread of heading directions differed significantly between Control and Cylinder conditions in cylinder position A and B, but not in C (Control vs Cylinder, k tests, p < 0.05 for all 3 distances in A and B, p > 0.05 for all 3 distances in C; Fig. 2). Similarly, overall path straightness was significantly decreased in A and B but not C (Index of straightness: Control vs. Cylinder, Mann Whitney U test, p < 0.001 in A, p < 0.05 in B, p > 0.05 in C). Average walking speed did not change in any of the conditions (Control vs. Cylinder, Mann Whitney U test, p > 0.05 in A, B and C). Also, in all groups, some ants returned back to the nest without leaving the arena, however, the proportion of ants returning to the nest did not differ between the Control and Cylinder conditions (Control vs. Cylinder, Chi-squared test, p > 0.05 for A, B and C). In summary, the presence of a novel cylinder in the nest vicinity induced learning walks when it was placed along the learnt route to the feeder (position A and B) but not when it was placed at position C.

**Figure 2:**
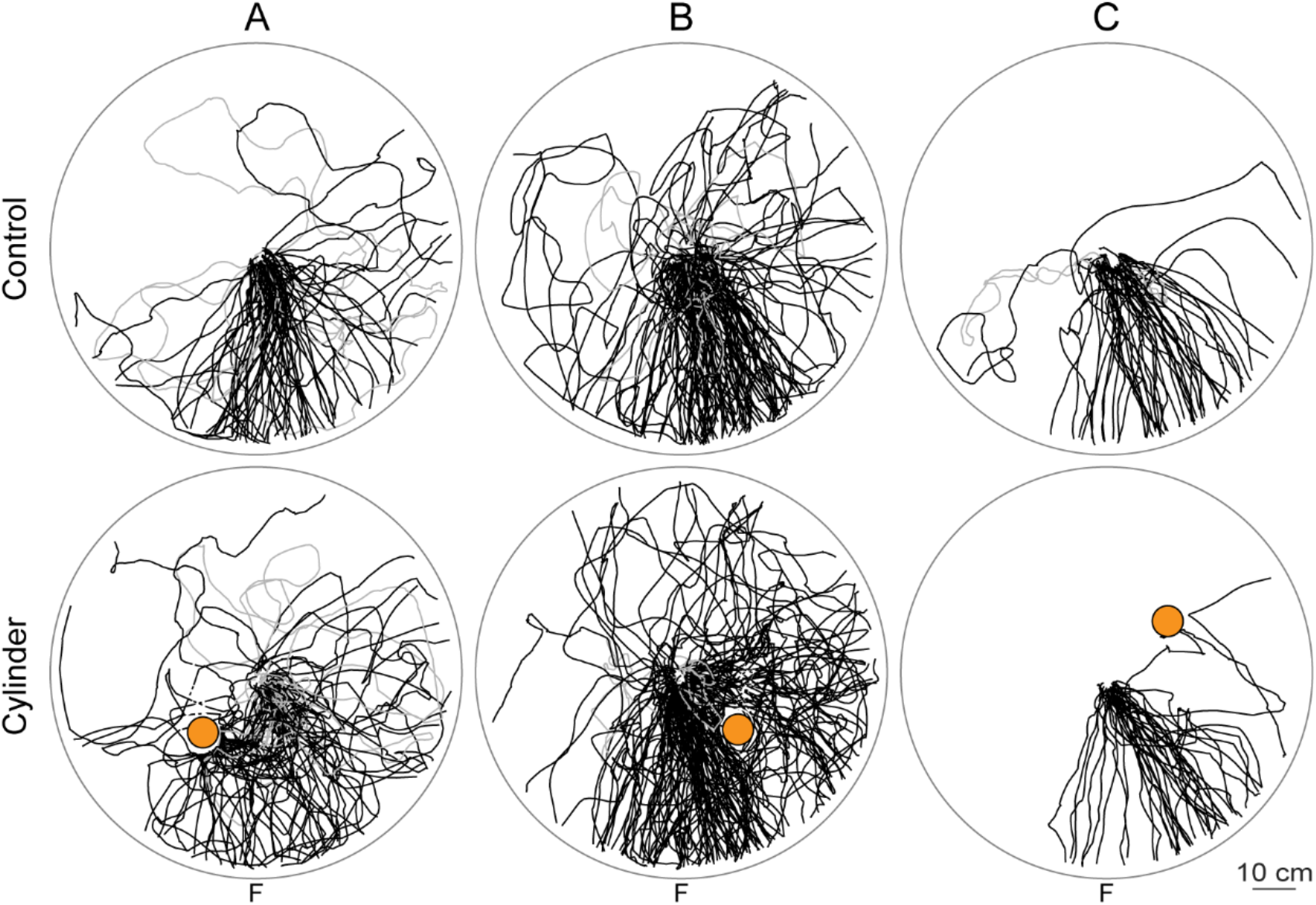
Outwards ants heading to the feeder when familiar with the nest surrounding (top) and when a novel cylinder is added (bottom). Trajectories from individual ants leaving the nest and entirely crossing the arena (1 m in diameter) are shown as black lines. Ants returning to the nest without leaving the arena are shown in grey. Dashed lines connect gaps in paths. Cylinder is shown in orange. Ants from different nests, but with the same cylinder positions (A, B or C) are grouped together. Prevailing wind on these days was 55° positive to the feeder direction. **A**: Jul-13, Jul-26, Jul-28; Control, n = 71 ants, nest returns (ants not entirely crossing the arena but returning to the nest): 5 out of 71 ants; Cylinder, n = 79 ants, nest returns: 12 out of 79 ants. **B**: Jul-07, Jul-14, Jul-16; Control, n = 129 ants, nest returns: 8 out of 129; Cylinder, n = 158 ants, nest returns: 15 out of 158 ants. **C**: Jul-17, Jul-27; Control, n = 53 ants, nest returns: 2 out of 53 ants; Cylinder, n = 39 ants, nest returns: 0 out of 39 ants.

When time allowed, we additionally recorded the ants’ first nest return after having added the cylinder (Fig. S1). Similar to what we have seen in outwards ants, we observed in inwards ants that the overall change in paths was bigger for cylinder position A than C (Fig. S1). However, Control ants from the cylinder position B were noisier than usual and thus made a comparison difficult.

### Structure of learning walks is shaped by the wind direction

To analyse how the fine structure of learning walks was influenced by the direction of the wind, we analysed in fine detail the ants’ walking speed (Fig. 3), turning behaviour (Fig. 4) and facing direction (Fig. 5) for cylinder positions A and B. In some conditions, the directional scatter of path chunks with low walking speed (Rayleigh test for circular means calculated for each ant: Control, Ai: p < 0.01, Aii: p > 0.05, Bi: p < 0.001, Bii: p < 0.001; Cylinder, Ai: p < 0.001, Aii: p > 0.05, Bi: p < 0.001, Bii: p < 0.05), turns (Rayleigh test for circular means calculated for each ant: Control, Ai: p > 0.05, Aii: p > 0.05, Bi: p > 0.05, Bii: p > 0.05; Cylinder, Ai: p > 0.05, Aii: p > 0.05, Bi: p < 0.05, Bii: p < 0.05) and nest alignments (Rayleigh test for circular means calculated for each ant: Control, Ai: p > 0.05, Aii: p > 0.05, Bi: p > 0.05, Bii: p > 0.05; Cylinder, Ai: p > 0.05, Aii: p > 0.05, Bi: p < 0.05, Bii: p < 0.05) was high, hence, we focused on cylinder position B (Bi: wind +55° to the feeder vs. Bii: wind −31° to the feeder). We found that the wind direction had a significant influence on the spatial distribution of path sections with low walking speed, turns and nest alignments, respectively. The spatial distribution of the path segments with low walking speed (Bi vs. Bii in Fig. 3; Watson Williams test, Control p < 0.01. Cylinder p < 0.001), turns (Bi vs. Bii in Fig. 4; Watson Williams test, Cylinder, p < 0.01) and also nest alignments (Bi vs. Bii in Fig. 5; Watson Williams test, Cylinder, p < 0.05) was significantly controlled by the direction of the wind. Hence, learning walks are adapted to take into account the current directionality of wind.

**Figure 3:**
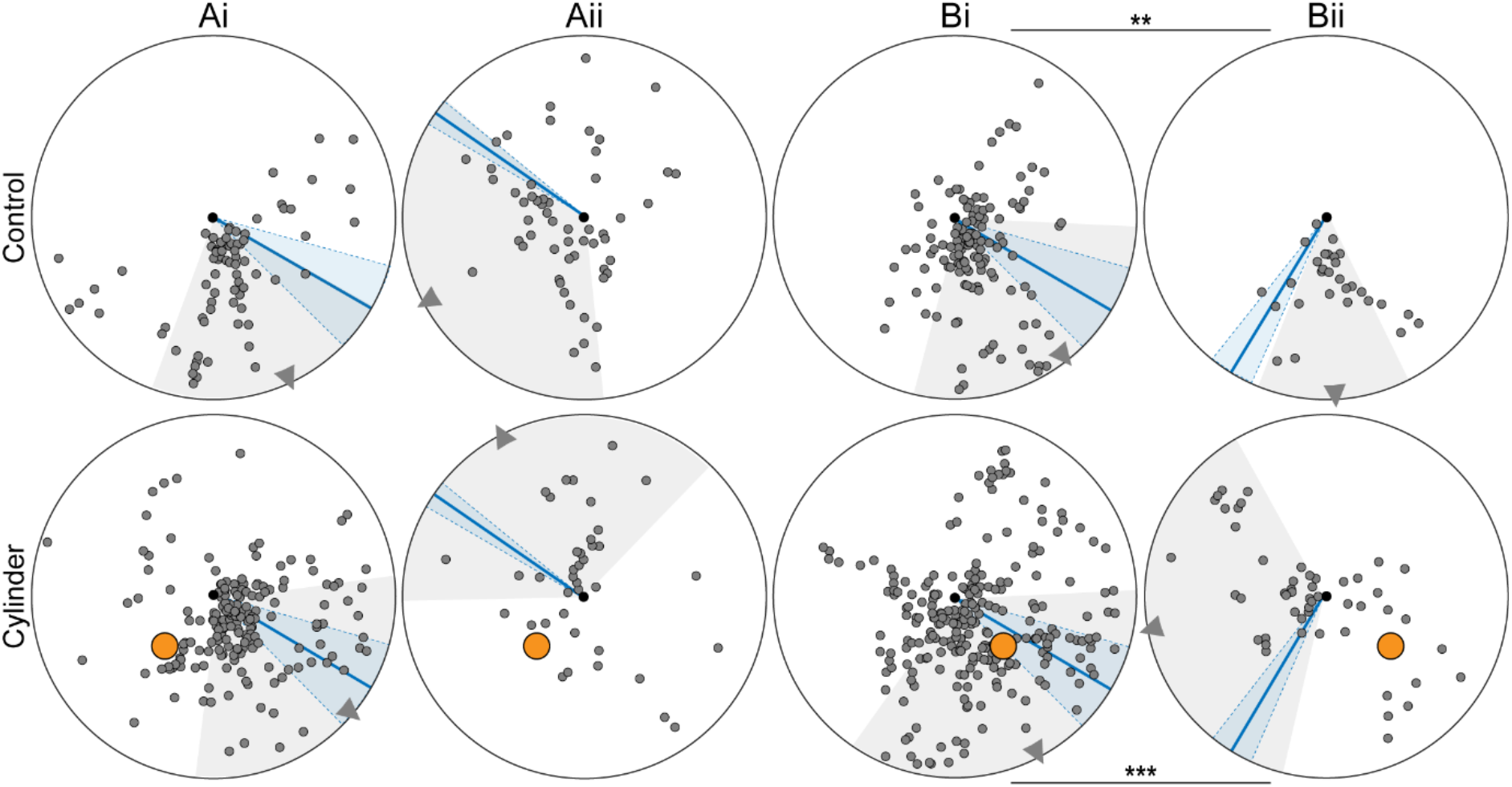
Path segments with low walking speed when ants are familiar with the nest surrounding (top) and when a novel cylinder is added (bottom). Path segments (2 cm chunks) where ants walked slower than 0.06 m/s are shown as small circles. Nest, black dot. Cylinder is shown in orange. Ants from different nests, but with the same cylinder position (A or B) are grouped together. Wind direction: mean with standard deviation (blue line and shadow). Ai and Bi: +55° to the feeder direction (prevailing wind), Aii: −125° to the feeder direction, Bii: −31° to the feeder direction. Direction of selected path chunks: mean with standard deviation (grey triangle and shadow). Watson Williams test, Bi vs Bii, Control p < 0.01 (**), Cylinder p < 0.001 (***).

**Figure 4:**
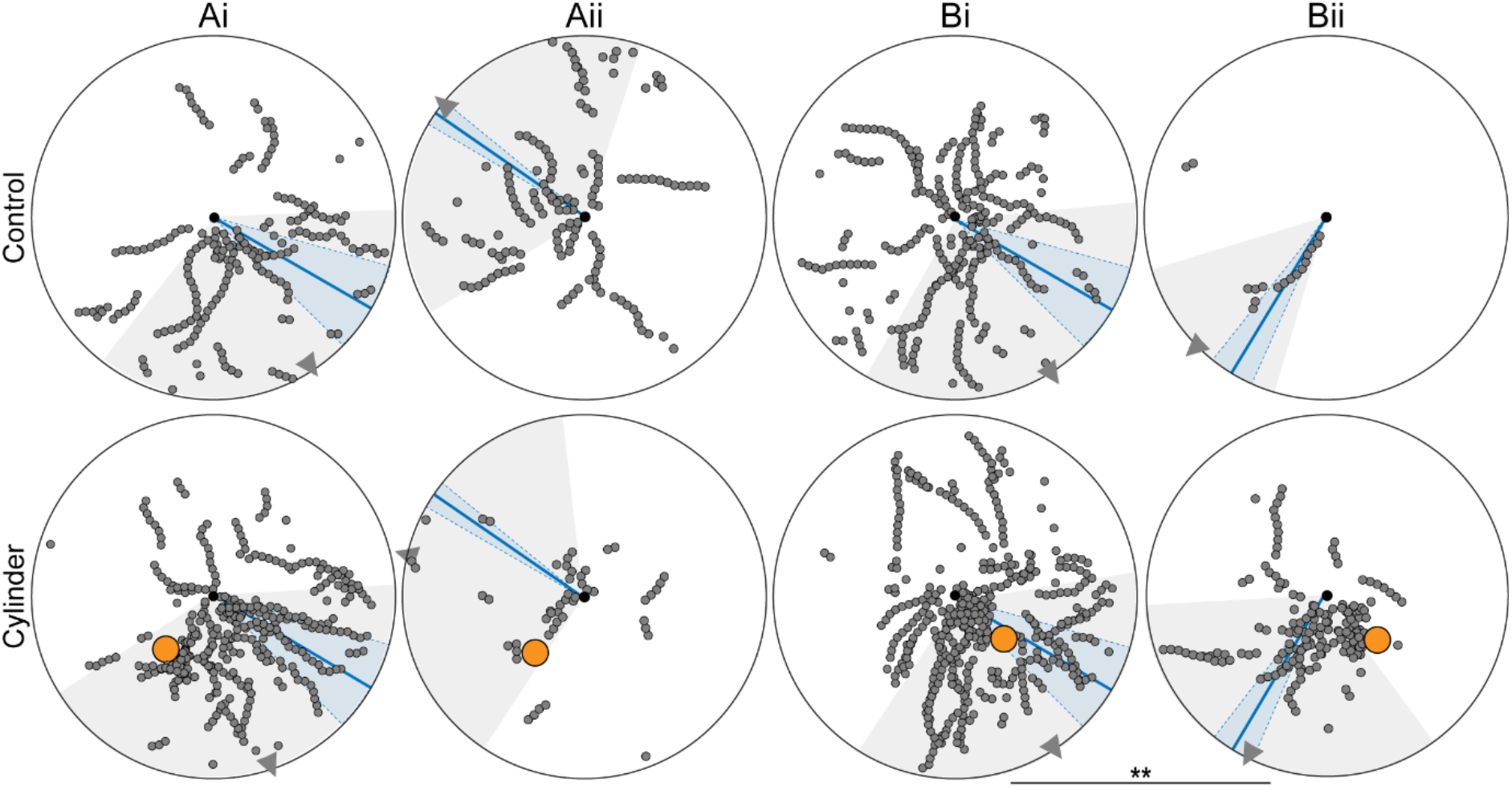
Path segments with turns when ants are familiar with the nest surrounding (top) and when a novel cylinder is added (bottom). Path chunks where ants turned are shown as small circles. Nest, black dot. Cylinder is shown in orange. Ants from different nests, but with the same cylinder position (A or B) are grouped together. Wind direction: mean with standard deviation (blue line and shadow). Ai and Bi: +55° to the feeder direction (prevailing wind), Aii: −125° to the feeder direction, Bii: −31° to the feeder direction. Direction of selected path chunks: mean with standard deviation (grey triangle and shadow). Watson Williams test, Bi vs Bii, Cylinder, p < 0.01 (**).

**Figure 5:**
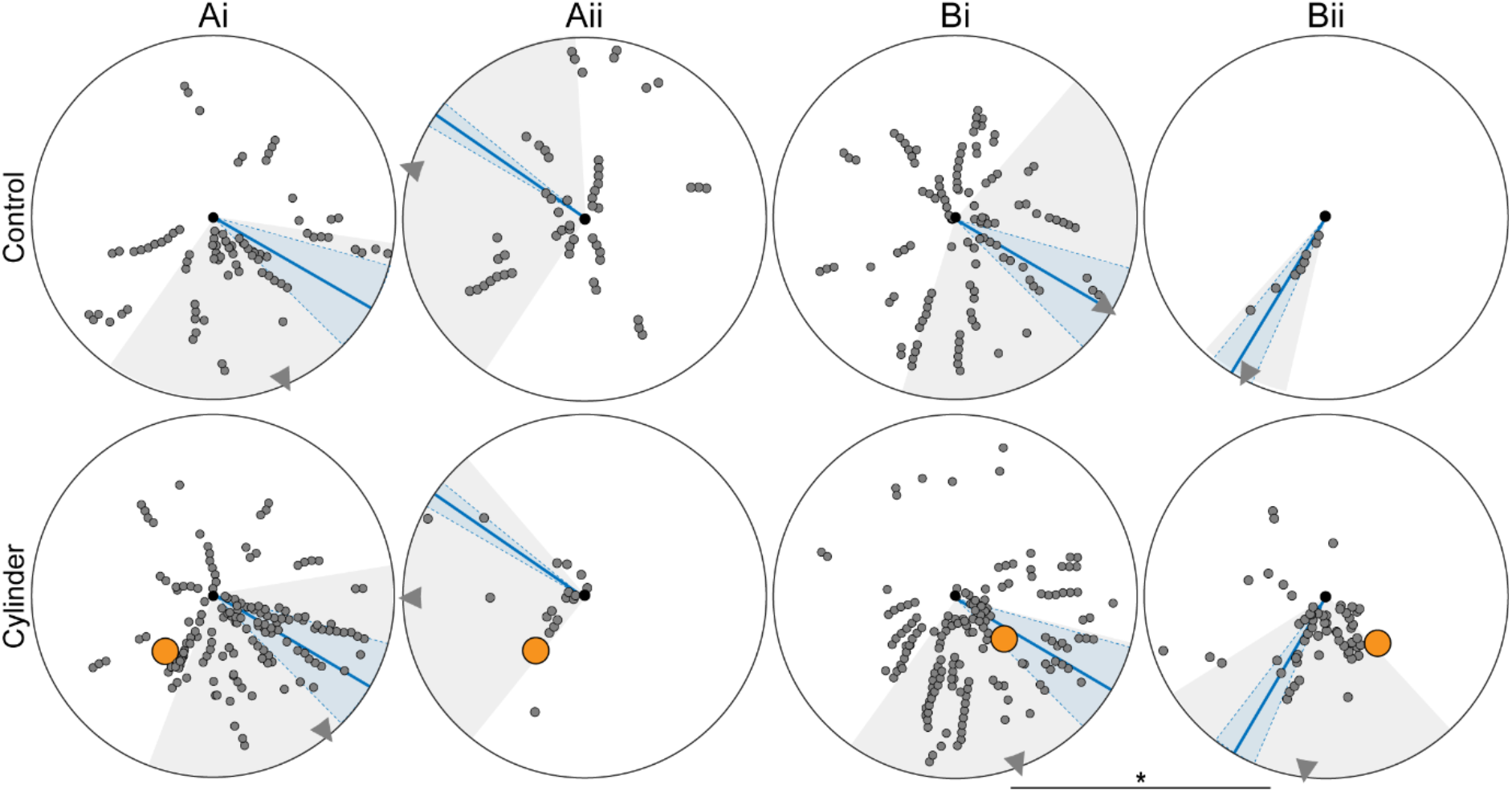
Nest alignments when familiar with the nest surrounding (top) and when a novel cylinder is added (bottom). Path segments that were aligned to the nest are shown as small circles. Nest, black dot. Cylinder is shown in orange. Ants from different nests, but with same cylinder positions (A or B) are grouped together. Wind direction: mean with standard deviation (blue line and shadow). Ai and Bi: +55° to the feeder direction (prevailing wind), Aii: −125° to the feeder direction, Bii: −31° to the feeder direction. Direction of selected path chunks: mean with standard deviation (grey triangle and shadow). Watson Williams test, Bi vs Bii, Cylinder, p < 0.05 (*).

In addition to the distribution, we looked at the proportion of ants that demonstrated the specific path parameters of interest. The proportion of ants with path segments of low walking speed did not differ between Control and Cylinder conditions (Control vs Cylinder, Chi-square test, all p > 0.05). However, ants more often performed turns and aligned themselves towards the nest, respectively, when the cylinder was added in condition Ai (Turns: Control vs Cylinder, Chi-square test, p < 0.001; Nest alignments: Control vs Cylinder, Chi-square test, p < 0.01) and Bii (Turns: Control vs Cylinder, Chi-square test, p < 0.01; Nest alignments: Control vs Cylinder, Chi-square test, p < 0.01), but not in Aii (Turns: Control vs Cylinder, Chi-square test, p > 0.05; Nest alignments: Control vs Cylinder, Chi-square test, p > 0.05) and Bi (Turns: Control vs Cylinder, Chi-square test, p > 0.05; Nest alignments: Control vs Cylinder, Chi-square test, p > 0.05).

## Discussion

Our aim was to investigate whether the structure of learning walks of *Cataglyphis fortis* desert ants takes into account wind direction in conjunction with the learning of new visual information. We trained foragers to visit a regular feeder and then introduced a black cylinder near their nest, thus learning walks (or re-learning walks) were provoked in regular foragers. By doing this across days with different prevailing wind directions, we were able to probe how ants compromise across information derived from different modalities.

It has been demonstrated previously that wind direction plays a role in the approach strategies of ants to unfamiliar (Buehlmann et al., 2014) and familiar feeding locations (Wolf and Wehner, 2000, 2005) as well as to the nest (Buehlmann et al., 2012). In all these scenarios the final approach is into the wind, which maximises the potential to use olfactory information. In agreement with these previous observations, we see that the departure directions of our ants are influenced by the current wind direction (Fig. 2), with forager routes deflected in such a way that ants will arrive downwind of their target. Additionally, by biasing the departure direction of ants’ foraging routes, the wind direction indirectly acts to determine the locations and structure of the provoked learning walks. From the perspective of an individual forager upon leaving the nest, the initial foraging direction is chosen as an integration of the remembered food direction, and the direction of the wind (as in (Wolf and Wehner, 2005)). If that path direction means that the ant’s path runs close by the newly installed landmark (A and B, vs C), then we see an upregulation in the sensori-motor motifs associated with learning. However, the landmark does not act as a direct driver of the precise locations of the learning behaviours. For instance, if the landmark position had a definitive influence on the structure of the learning walks, inspections following the landmark being added to positions A and B would be mirror-symmetric to each other, given the same wind conditions.

To indicate those portions of routes that might be indicative of ‘re-learning walks’, we looked for slowing (Fig. 3), turning (Fig. 4) and nest inspections (Fig. 5). We know that across a range of ant species there are significant differences in the sensori-motor motifs that signify learning walks (Fleischmann et al., 2017; Zeil and Fleischmann, 2019). The sensory ecology of an ant species seems to be the major determinant of the style of their learning walks, although most of our understanding comes from examples of ant species in habitats with different amounts of visual clutter. Our results suggest that other sensory modalities also need to be considered when cataloguing the relationship between learning walks and habitat. It might be that *Cataglyphis fortis* foragers take advantage of the relatively constant wind direction to enhance the utility of learning walks, and whilst they are clearly influenced by the addition of the landmark, they may place less weight on the precise learning of the visual cue. Further experiments, especially with brand new foragers, would allow us to investigate how individual species give appropriate weight to wind information relative to the information from visual clutter in their habitat.

In order to implement these experiments, we needed to record ants on days with a range of wind conditions, however it is true that for this habitat there was a prevailing wind direction which was seen on most days. The influence of this being that most foraging paths had a characteristic deflection, and so over a day-to-day period the habitual routes of foragers would have this shape. This raises the question of to what extent wind information is incorporated into the route memories of regular foragers. Does the wind set a route direction, but then route guidance information is learnt independently of the wind? Or, is wind information part of the multimodal sensory information used to guide the habitual routes? The first of these options is a pattern of route learning that is seen in other modalities (Collett et al., 2003) when a particular cue determines (or scaffolds) the shape of the learnt habitual route, but then, for experienced foragers, the habitual route can be accurately navigated without the original scaffold. This pattern of learning can be seen when visual cues (Graham et al., 2003), path integration (Collett et al., 2001) or pheromone trails (Harrison et al., 1989) are used to determine an initial path shape, which is still maintained in experienced foragers, even when the original scaffold is removed. In these experiments we did not have marked ants of known identity or experience, therefore it is impossible to determine the ontogeny of route development. However, on days when the wind direction showed a strong change away from the prevailing direction, we do see significant disturbance to outwards paths. Assuming that a significant proportion of the disturbed ants were experienced individuals, it suggests that wind cues are part of the multimodal suite of cues used to guide routes (Buehlmann et al., 2020), rather than simply a temporary scaffold.

## Acknowledgements

This project was funded by the people programme (Marie Curie Actions) of the European Union’s Seventh Framework Programme (FP7/2007-2013, under REA grant agreement no. PIEF-GA-2013-301624765) to CB. PG is additionally funded by a BBSRC grant BB/R005036/1 and EPSRC grant EP/P006094/1 and CB is additionally funded by a BBSRC grant BB/R005036/1. JAVV was hosted within the Brains on Board project.

## Compliance with ethical standards

### Ethical approval

All applicable international, national, and institutional guidelines for the care and use of animals were followed.

---

**Figure S1:**
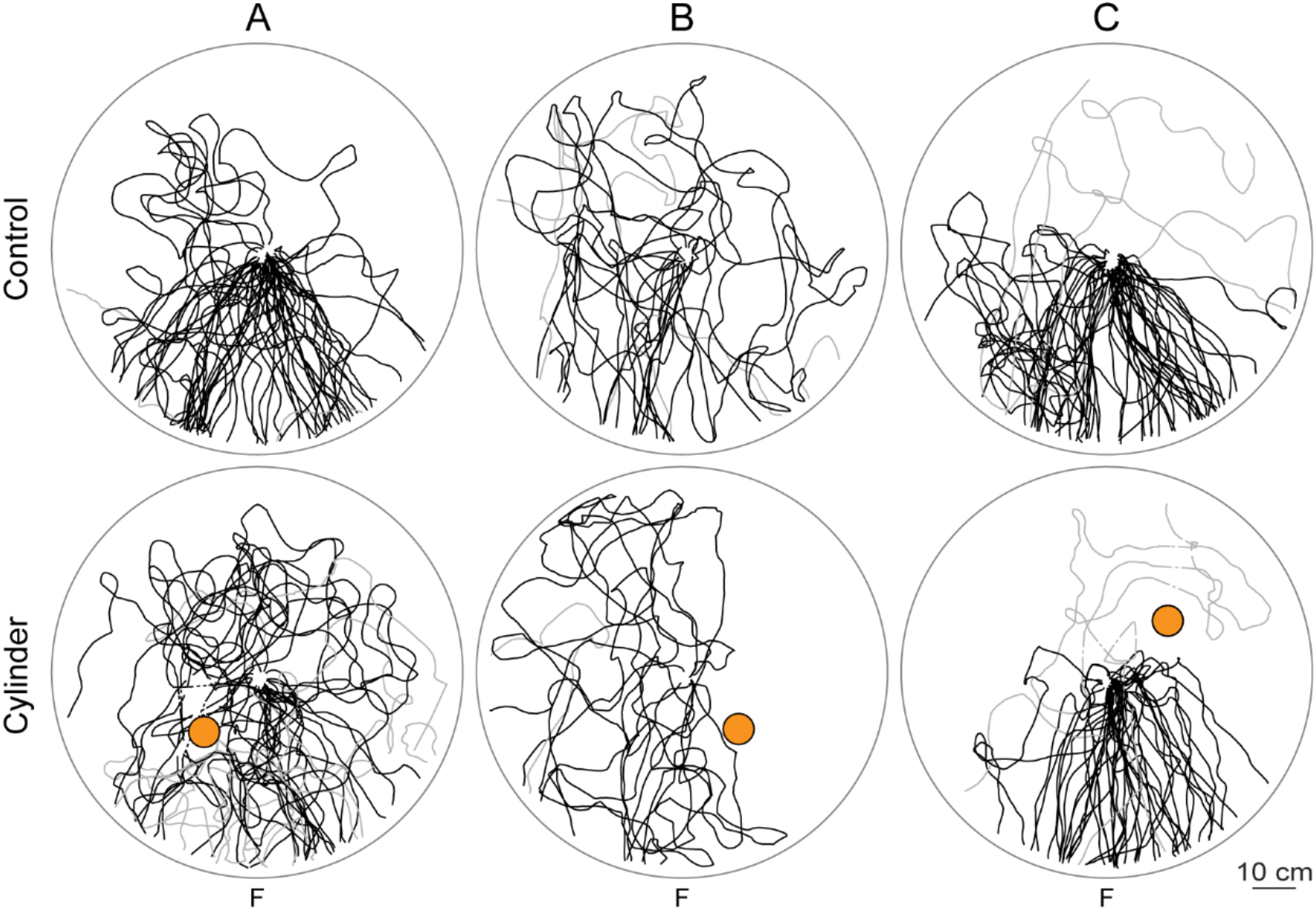
Inwards ants returning to the nest when familiar with the nest surrounding (top) and when an unfamiliar cylinder is added (bottom). Trajectories from individual ants coming from the feeder and entering the nest (1 m in diameter) are shown as black lines. To focus on ants coming from the feeder, only ants coming from the feeder direction +/- 90° were considered here. Ants crossing the arena without entering the nest are shown in grey. Dashed lines connect gaps in paths. Cylinder is shown in orange. Ants from different nests, but with the same cylinder positions (A, B or C) are grouped together. Prevailing wind on these days was 55° positive to the feeder direction. **A**: Jul-13, Jul-26; Control, n = 56 ants, not entering nest: 2 out of 56 ants; Cylinder, n = 35 ants, not entering nest: 7 out of 35 ants. **B**: Jul-07; Control, n = 24 ants, not entering nest: 4 out of 24; Cylinder, n = 11 ants, not entering nest: 1 out of 11 ants. **C**: Jul-27; Control, n = 55 ants, not entering nest: 4 out of 55 ants; Cylinder, n = 48 ants, not entering nest: 3 out of 48 ants. The number of ants not entering the nest but leaving the arena again was increased for cylinder position A but not for B and C (Number of ants not entering the nest: Control vs. Cylinder, Fisher’s exact tests, p < 0.05 in A, p > 0.05 in B and C). Similarly, path straightness was decreased in A but not in B and C (Index of straightness: Control vs. Cylinder, Mann Whitney U test, p < 0.001 in A, p > 0.05 in B and C). Furthermore, we found differences in walking speed (Walking speed: Control vs. Cylinder, Mann Whitney U test, p > 0.05 in A and C, p < 0.01 in B).

